# Reduced interhemispheric and increased intrahemispheric connectivity in schizophrenia brain networks

**DOI:** 10.1101/755173

**Authors:** Jorne Laton, Jeroen Van Schependom, Jeroen Decoster, Tim Moons, Marc De Hert, Guy Nagels, Maarten De Vos

**Affiliations:** Institute of Biomedical Engineering, Department of Engineering Science, University of Oxford Old Road Campus Research Building, Headington, Oxford OX3 7DQ, United Kingdom; Center for Neurosciences (C4N), UZ Brussel, Vrije Universiteit Brussel Pleinlaan 2, 1050 Brussel, Belgium; UPC KU Leuven - Campus Kortenberg, Department of Neurosciences, KU Leuven Leuvensesteenweg 517, 3070 Kortenberg, Belgium

**Keywords:** resting state, oddball, mismatch, coherence, directed transfer function, corpus callosum

## Abstract

**Introduction:** Brain connectivity is disturbed in schizophrenia, both during resting state and during active tasks. Schizophrenia is characterised by a corpus callosum pathology and an inability to suppress overstimulation, both of which relate to this disturbed connectivity. We wanted to verify whether network analysis on EEG sensor level can reveal the corpus callosum pathology in schizophrenia.

**Methods:** We measured 62-channel EEG on 46 schizophrenia patients and 43 healthy controls during eyes-closed and eyes-open resting-state, mismatch negativity and visual and auditory oddball. We assessed connectivity through correlation, coherence and directed transfer function (DTF) in the delta, theta, alpha, low- and high beta bands.

**Results:** The coherence and the DTF picked up a consistent pattern of reduced interhemispheric and enhanced intrahemispheric connectivity strength in schizophrenia in the alpha and beta band. This disturbance pattern appeared across all paradigms in the parietal and the occipital region and was generally more pronounced in the right hemisphere.

**Conclusions:** This is the first study to use multiple similarity measures and different tasks to confirm disturbed brain connectivity on EEG sensor level. We hypothesise that the interhemispheric reductions reflect transcallosal disconnection, while the intrahemispheric increases indicate the inability to suppress the response to stimuli.

## 1. Introduction

Schizophrenia is a heritable chronic brain disorder affecting 1 out of 100 people (Lewis and Lieberman, 2000; Javitt and Sweet, 2015). Symptoms generally arise between the ages of 15 to 35 years, while the average life expectancy is 12 to 15 years less than for people without brain disorder. This is partly due to a higher suicide rate of 10 % for people with schizophrenia (Lewis and Lieberman, 2000). The classical diagnosis of the disease is performed by an experienced psychiatrist, who bases his diagnosis on the outward symptoms. These are divided in three groups: positive, negative and cognitive symptoms (Lewis and Lieberman, 2000). Positive symptoms are thoughts, behaviours, or sensory perceptions that normal people generally do not experience. In contrast, negative symptoms correspond to an absence or diminution sensation, which is considered normal. Cognitive symptoms are related to troubles with memory function and concentration (Javitt and Sweet, 2015). The individual combination of these outward symptoms is the exophenotype of a patient.

In contrast, the endophenotype is the individual combination of internal signs of the disease. Endophenotypes are useful to differentiate people with schizophrenia in greater detail, in particular the neurophysiological endophenotypes, which have traditionally relied on direct assessment of the activity intensity, usually by examination of amplitude differences in a peak analysis. Deficits in the auditory system have been associated with schizophrenia (Pfefferbaum *et al.*, 1989; Polich, 2007; Roach and Mathalon, 2008; Mathalon *et al.*, 2010; Decoster *et al.*, 2012; Näätänen *et al.*, 2012; Nagai *et al.*, 2013; Bartha-Doering *et al.*, 2015; Thibaut *et al.*, 2015; Javitt and Sweet, 2015; Schall, 2016). Differences in cognitive processing have been revealed in visual paradigms too (Bestelmeyer, 2012; Oribe *et al.*, 2014; Farkas *et al.*, 2015), as well as during resting state (Greicius, 2008; Alonso-solís *et al.*, 2015; Di Lorenzo *et al.*, 2015). Quantifying the EEG amplitude during attentional tasks even allowed to identify patients with schizophrenia versus controls in 84.7 % of cases (Laton *et al.*, 2014). The additional information provided by these endophenotypes may also facilitate the development of therapies targeting the specific underlying mechanisms causing schizophrenia (Huang, Chen and Zhang, 2015; Rosen, Spellman and Gordon, 2015).

More recently, network analysis to assess brain connectivity rather than activity has gained importance in neurophysiological research (Fornito and Bullmore, 2015). Several studies have shown that brain connectivity is also disturbed in schizophrenia patients during resting state (Andreou *et al.*, 2014, 2015; Unschuld *et al.*, 2014; Alonso-solís *et al.*, 2015; Olejarczyk and Jernajczyk, 2017; Larsen *et al.*, 2018) and in individual task-related EEG measurements such as the auditory oddball (Bachiller *et al.*, 2015).

Network analysis of neurophysiological signals consists of determining the similarity and/or the causality between the signals measured at different locations to estimate the real underlying connectivity. Altered similarity or causality can be an indicator of disturbed physical links between brain regions, or the appearance of new links. It can also tell something about regions you cannot directly measure, such as the corpus callosum, through which a lot of communication happens in the brain. Examination of similarity measures was used to demonstrate decreased transcallosal connectivity in schizophrenia (Crow, 1998; Foong *et al.*, 2000; Whitford *et al.*, 2010).

In this study, we examined the endophenotype of schizophrenia through the active functional networks in resting state, two auditory paradigms and a visual paradigm. We investigated differences between adjacency networks calculated on schizophrenia patients and on healthy controls. We used three different edge detection methods to quantify differences in functional networks: correlation, coherence and directed transfer function (DTF). The reason for using multiple methods, is that all of these rely on certain assumptions and we want to assess which network patterns are picked up systematically. The DTF quantifies directed functional connectivity without being affected by volume conduction effects. It is derived from the multivariate model coefficients generated by estimating spectral Granger causality between electrodes (Kaminski and Blinowska, 2014). To enable an easy comparison with literature, we also included the correlation and the coherence in the standard frequency bands. Correlation is a simple measure quantifying linear relationships, but does not give information on the spectral properties of the signal and is sensitive to volume conduction. Coherence can be seen as correlation in the spectral domain, allowing splitting up the networks in different bands, but it is also sensitive to volume conduction.

To our knowledge, this is the first study that focusses on rest and attention-related connectivity in a battery of multimodal attention-related tasks in schizophrenia patients. As schizophrenia is a very heterogeneous disease, it is important to include a rather broad spectrum of activities, ranging from passive (resting state) and semi-active (mismatch negativity) to active (oddball) and invoking visual and auditory functions. We want to verify whether network analysis on EEG sensor level can reveal the corpus callosum pathology in schizophrenia (Crow, 1998; Foong *et al.*, 2000; Whitford *et al.*, 2010).

## 2. Methods

### 2.1 Participants

We included 43 healthy non-medicated controls and 46 patients with schizophrenia or schizoaffective disorder who underwent EEG during the auditory (AO) and the visual oddball (VO) paradigm, the mismatch negativity (MN) paradigm and resting state (RS) with eyes open and closed. Patients were recruited at the Universitair Psychiatrisch Centrum (UPC) KULeuven – Campus Kortenberg, where they were diagnosed with a semi-structured interview (OPCRIT v4.0). The methods were carried out in accordance with the relevant guidelines and all experimental protocols were approved by the ethical review board of the UPC KULeuven – Campus Kortenberg. All participants have given written informed consent.

### 2.2 Recording conditions

The EEG equipment at the hospital consisted of a 64-channel ANT digital measure station (ANT, The Netherlands). Ag-AgCl electrodes were arranged in an electrode cap using the international 10-10 system (Nuwer *et al.*, 1998). Signals were digitised at a sampling frequency of 256 Hz and stored for offline analysis.

### 2.3 Paradigms

#### 2.3.1 Resting state

Participants were sitting at rest, first with their eyes closed for four minutes and then four minutes with their eyes open. During eyes open, they were asked to focus on a fixation point and during eyes closed, they had to try moving their eyes as little as possible.

For the resting-state analysis, we omitted six patients, five of whom had no rest data and one with a flat channel.

#### 2.3.2 Auditory oddball

This paradigm consisted of target, distractor and standard tones. The target and the distractor differed from the standard tone (1000 Hz) respectively by a higher (1500 Hz) and a lower frequency (500 Hz). All had a duration of 100 ms and a loudness of 70 dB. These stimuli were presented pseudo-randomly, with a distribution of 80% standard, 10% target and 10% distractor and an interstimulus interval randomised between 1 and 1.5 seconds. There were 400 stimuli per test, resulting in a total test time of 540 seconds.

#### 2.3.3 Visual oddball

For the visual P300, a target (square, side 106 pixels), distractor (circle, diameter 176 pixels) and standard figure (square, side 158 pixels) were presented in full blue (RGB = 0, 0, 255) in the centre of a black (RGB = 0, 0, 0) background with a resolution of 1024 by 768 pixels. The distribution of the stimuli and the total test time were the same as for the auditory oddball paradigm.

#### 2.3.4 Mismatch negativity

The paradigm consisted of duration-deviant, frequency-deviant and standard tones. The duration- and frequency deviant tones differed from the standard tone (100 ms, 1000 Hz) respectively by a longer duration (250 ms) and a higher frequency (1500 Hz) and all have a loudness of 70 dB. These stimuli were presented in pseudo-random order, with a distribution of 90% standard, 5% duration deviant and 5% frequency deviant and an inter-stimulus interval of 300 ms. There were 1800 stimuli per test, resulting in a total test time of 733 seconds.

### 2.4 Pre-processing

Filtering was done in Matlab (MATLAB, 2019) using SPM12 (Ashburner *et al.*, 2014). Three Butterworth filters were applied: a high-pass filter with a cut-off frequency at 0.1 Hz, removing DC, a low-pass filter with a cut-off frequency at 30 Hz and a band-stop filter with range of 48 Hz to 52 Hz, removing a remaining 50 Hz mains hum.

After this, pre-processing was ported to EEGLab 14.1.2 (Delorme and Makeig, 2004). Data were rereferenced to average (Dien, 1998), followed by reference electrode standardisation (REST) (Yao, 2001; Dong *et al.*, 2017), using electrode positions estimated by the BESA four-shell spherical model. REST reduces volume conduction and reference bias and is recommended for network analysis, especially in high-density setups with more than 60 (preferably 100) electrodes (Chella *et al.*, 2016; Olejarczyk and Jernajczyk, 2017).

We extracted and baseline-corrected one-second epochs (baseline: 200ms) in the oddball paradigms, 600ms epochs (baseline: 100ms) in the mismatch negativity and four-second epochs in the resting-state data. We ran automatic artefact rejection to remove large fluctuations and improbable activity (Delorme and Makeig, 2004).

### 2.5 Edge detection

Network edges were calculated between each pair of channels and stored in an adjacency matrix. We investigated correlation, coherence and directed transfer function. These are all normalised measures for which a higher value means a stronger relation or edge between a pair of electrodes.

#### 2.5.1 Pearson Correlation

This is the simplest and most straightforward measure of similarity between two time signals. Its limitations are that it only shows linear relations, it gives no spectral information and it is sensitive to volume conduction and signal amplitude.

#### 2.5.2 Coherence

This can be seen as a correlation in the frequency domain and is the absolute value of the normalised cross spectrum, or coherency,

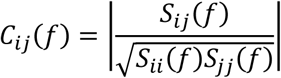

with *S*_*ij*_(*f*) the cross-spectrum between channels i and j.

Limitations of coherence are also sensitivity to volume conduction and signal amplitude (Van Schependom *et al.*, 2014).

We based our implementation of the coherence on the HERMES toolbox (Niso *et al.*, 2013). We concatenated the epochs into one signal and used a non-overlapping sliding window that was exactly aligned with the epochs, omitting the baseline. In the oddball paradigms, we therefore had 800ms windows (205 time points), in the mismatch paradigm, we had 500ms windows (128 time points) and in the resting state, windows were 4 seconds long (1024 time points). The number of windows was equal to the number of epochs: on average 39 for oddball, 77 for mismatch and 57 for resting state. There was no significant difference in the number of epochs between the groups for any of the paradigms or stimuli. We summarised the coherence in five separate bands by taking the average over each band: delta (1-4 Hz), theta (4-8 Hz), alpha (8-12 Hz), low beta (12-20 Hz) and high beta (20-30 Hz).

#### 2.5.3 Directed Transfer Function

The directed transfer function (DTF) does not have the limitations of the correlation and coherence, but assumes that the data can be approximated by an autoregressive model. It could have been called *directed coherence*, but *directed transfer function* was considered a more appropriate term (Kaminski and Blinowska, 1991). It is based on the principle of Granger Causality:

Given *X*(*t*), described by the following multivariate process:

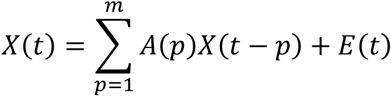

Transformed into the frequency domain, this is

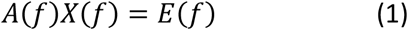

In which

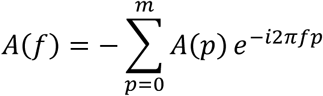

and *A*(0) = −I. Equation (1) can be rewritten to

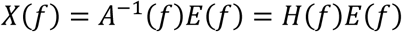

H is the transfer matrix of the system. The normalised DTF is defined as

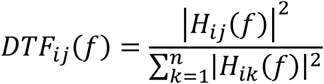

(Kaminski and Blinowska, 1991; Kaminski *et al.*, 2001).

An inconvenience of the DTF is that the number of data points in one window (*N* − *p*) needs to be at least an order of magnitude larger than *k* × *p*, with *k* the number of channels and *p* the model order (Blinowska, 2011):

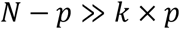

We are dealing with event-related potentials, of which a single trial is not long enough to fulfil the above condition. However, we can see the collection of ERPs from trials as an ensemble of realisations of a non-stationary stochastic process with locally stationary segments (Ding *et al.*, 2000; Pereda, Quiroga and Bhattacharya, 2005; Niso *et al.*, 2013). Therefore:

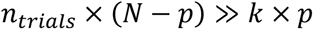

Using the Schwarz Bayesian Criterion and the Akaike Information Criterion, we found the optimal model order *p* = 4. With *N* = 205 (omitting the baseline) and *n*_*trials*_ ≥ 16 for the oddball paradigms, *N* = 128 and *n*_*trials*_ ≥ 39 for the mismatch paradigm and *k* = 62, this condition was met for every participant by at least a factor 10.

We summarised the DTF in the same bands as the coherence.

### 2.6 Group analysis

#### 2.6.1 Effect size

Cohen’s d, the effect size used to indicate standardised difference, was calculated between patients and controls for each of the modalities on adjacency matrices averaged per participant, using the controls as the reference.

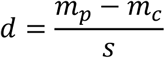

with *m*_*p*_ and *m*_*c*_ the patient and control averages, respectively, and *s* the pooled standard deviation.

#### 2.6.2 Correction for multiple comparisons

Student’s t-test was Bonferroni-corrected for multiple comparisons. This is the most conservative correction and likely results in an underestimation of the number of significant results. However, this provides a high certainty on the significant differences that are found. We did not include the number of bands nor the number of paradigms in this correction, as this would make the significance threshold too stringent, generating too many false negatives. Across bands, we only looked at the band which showed the largest number of differences and discarded non-overlapping differences found in the other bands. Across paradigms, we only looked at recurring difference patterns and discarded other differences that were not part of these patterns.

The number of unique directed channel pairs was

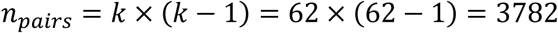

For the directed transfer function, this puts the threshold for significance to

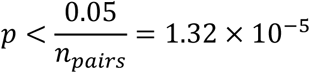

Correlation and coherence are symmetrical, so the significance threshold is

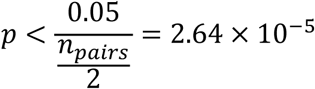

## 3. Results

### 3.1 Visualisation

The figures in the top row show the group averages in channel POz of the response to each of the stimuli of the paradigm. The healthy control group is plotted in blue and the schizophrenia group in red. The time after stimulus in milliseconds is on the horizontal axis, and the intensity in microvolts is on the vertical axis. The latencies and amplitudes of the most prominent peaks detected in POz are shown in table 1. For the oddball, this peak was the P3 and for the mismatch, this was the N2.

**Table 1:** Peaks in POz. Latencies are in ms and amplitudes in microvolts.

|  |  |  |  | HCT |  | SCH |  | p |
| --- | --- | --- | --- | --- | --- | --- | --- | --- |
|  |  |  |  | Mean | Std | Mean | Std |  |
| Auditory Oddball | Distractor | Latency | P3 | 384.1 | 61.2 | 406.1 | 91.7 | 0.190 |
|  |  | Amplitude | P3 | 8.6 | 4.5 | 8.0 | 6.5 | 0.606 |
|  | Target | Latency | P3 | 402.7 | 70.8 | 444.4 | 89.8 | 0.018 |
|  |  | Amplitude | P3 | 12.4 | 5.7 | 11.0 | 7.3 | 0.319 |
| Visual Oddball | Distractor | Latency | P3 | 404.2 | 73.5 | 420.8 | 81.0 | 0.315 |
|  |  | Amplitude | P3 | 4.9 | 3.2 | 3.9 | 3.2 | 0.141 |
|  | Target | Latency | P3 | 490.6 | 54.4 | 490.1 | 77.3 | 0.972 |
|  |  | Amplitude | P3 | 11.7 | 7.1 | 8.2 | 5.4 | 0.010 |
| Mismatch | Duration | Latency | N2 | 321.9 | 31.3 | 336.6 | 39.6 | 0.057 |
|  |  | Amplitude | N2 | -1.3 | 2.0 | -1.2 | 1.5 | 0.939 |
|  | Frequency | Latency | N2 | 229.1 | 44.3 | 232.1 | 44.2 | 0.752 |
|  |  | Amplitude | N2 | -1.4 | 1.7 | -1.4 | 1.5 | 0.965 |

The figures in the middle and bottom rows show network plots, respectively for the coherence and for the directed transfer function. A red line between two electrodes indicates a significant similarity increase in the schizophrenia group compared to the healthy control group. Likewise, blue lines show decreases. The plots are in top view, with the nose pointing north.

We visualised the direction of the DTF by an arrow in the direction of the information flow: from the influencing electrode towards the influenced electrode. A red arrow indicates an increased influence originating from the starting electrode.

### 3.2 Resting state

The correlation, the delta and theta coherence and the alpha and beta DTF were not able to pick up any group differences. In the alpha, low- and high beta band, we found a clear pattern of intrahemispheric increases – most strongly in the right hemisphere - and interhemispheric coherence decreases in the parietal region. This pattern was most apparent in the low beta band, which is shown in the top row of figure 1. In delta DTF, many connections were weaker in the schizophrenia group, mainly the ones of which the influencing electrode was centrally to parietally located (left and right). This was mainly in electrode pairs where the influencing electrode was parietal and close to the midline. As opposed to the coherence, these decreases were both inter- and intrahemispheric (bottom row in figure 1). The intrahemispheric DTF decreases generally had a longer range than the intrahemispheric coherence increases, which could be related to the band. In the eyes-open condition, we also saw increased intrahemispheric DTF values for connections originating from the left-temporal region towards left-frontal and left-parietal regions.

**Figure 1:**
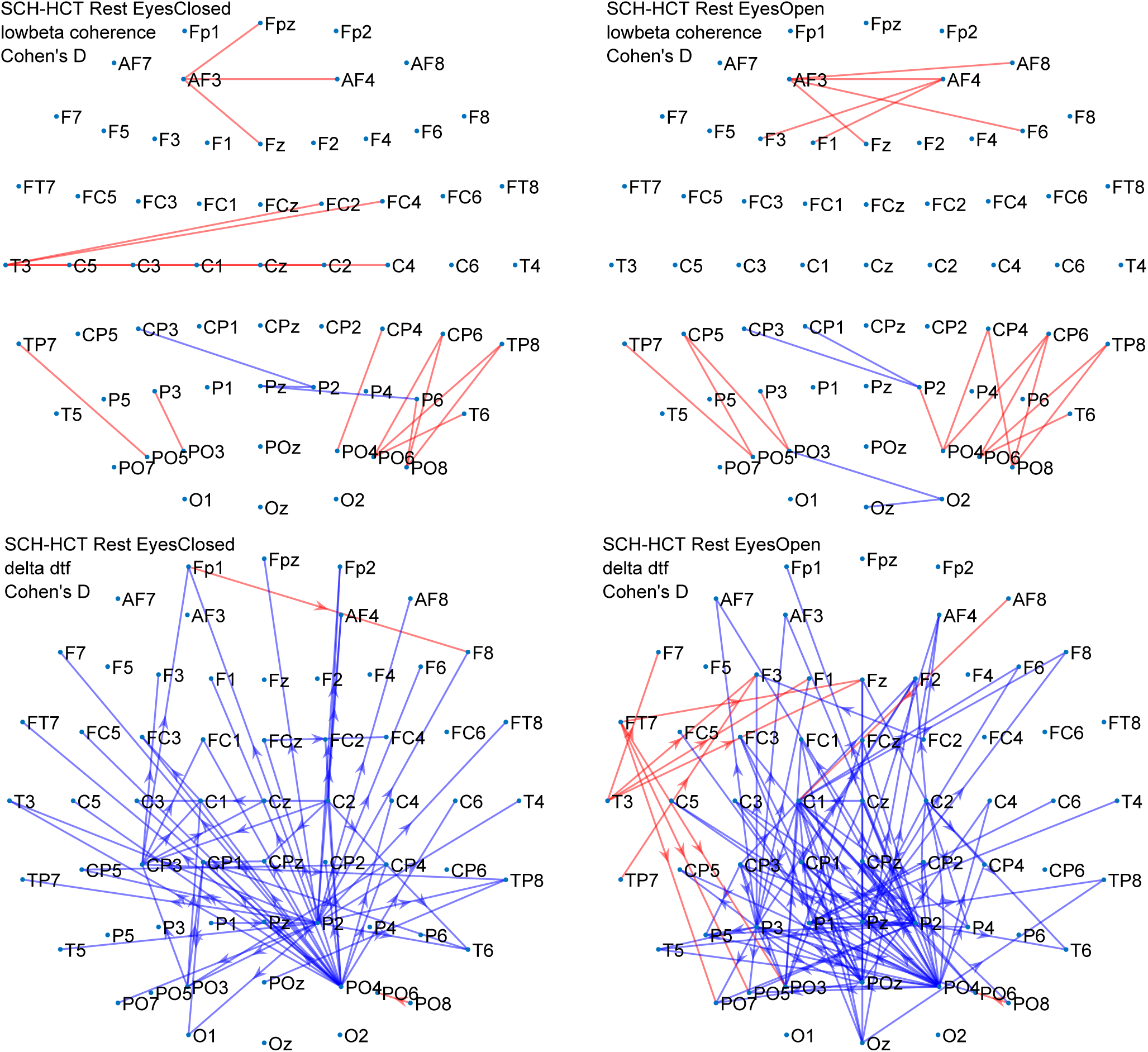
Rest eyes closed and -open coherence and DTF differences.

### 3.3 Auditory oddball

The correlation, the delta and theta coherence and the delta, theta and low-beta DTF did not pick up group differences, while in alpha and beta again a coherence pattern of interhemispheric decreases and intrahemispheric increases was found (middle row of figure 2). This pattern was situated in the parietal and occipital region - with increases mainly in the right hemisphere - and most prominent in the low beta band. It was characterised by a larger amount of interhemispheric decreases than the resting state, but mainly in the target response. In the DTF, there were only a couple of right-parietal electrodes of which the influence was significantly different. The very few significant differences did not seem consistent with the coherence, although long-range DTF decreases were all interhemispheric (bottom row in figure 2).

**Figure 2:**
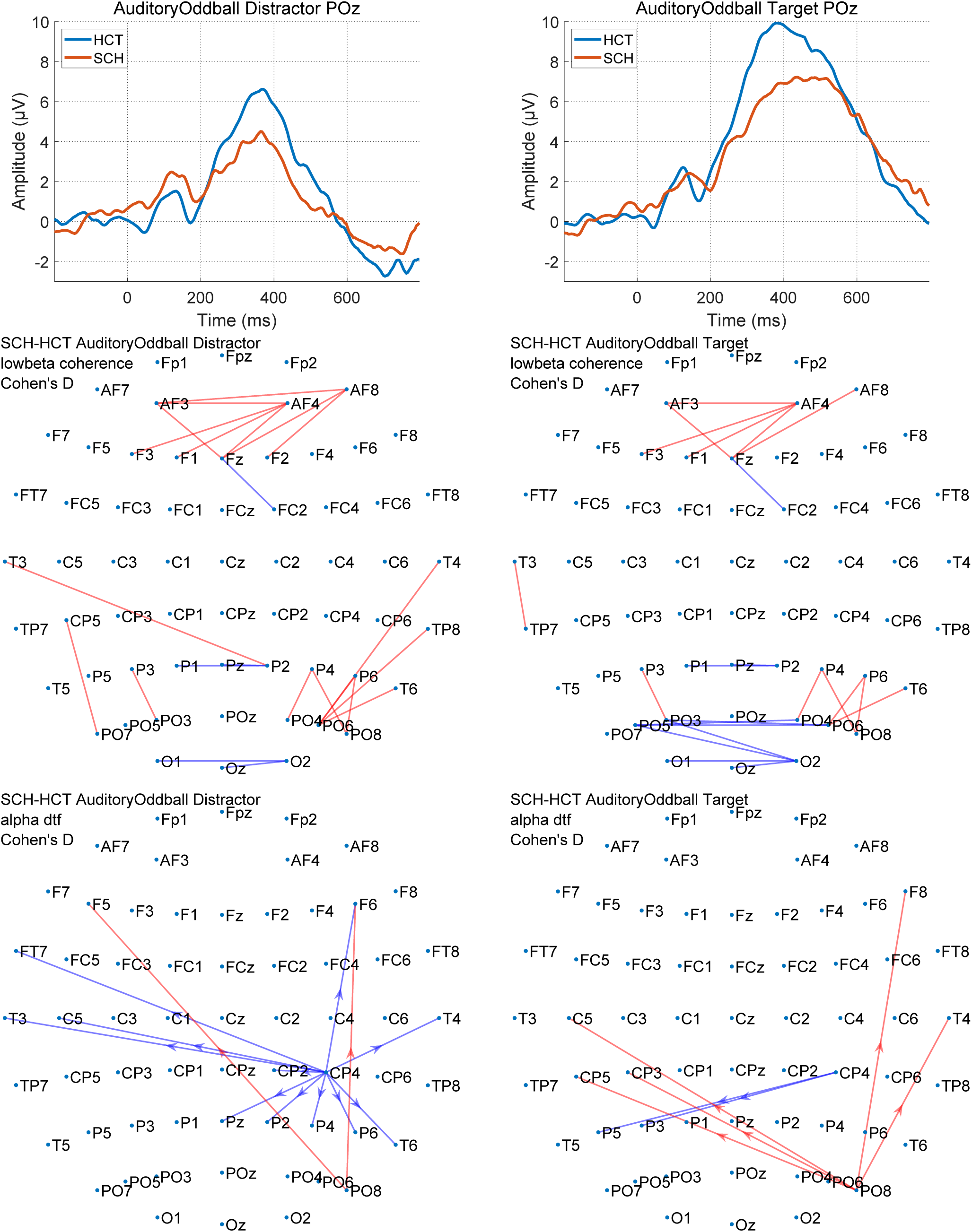
Auditory oddball distractor and target.

### 3.4 Visual oddball

The correlation, the delta and high beta coherence and the delta, theta and low-beta DTF did not pick up any differences. We saw the same parietal-occipital pattern as in the auditory oddball, but now in theta, alpha and low beta and with more intrahemispheric increases, particularly in the right hemisphere (middle row in figure 3). Similar to the auditory oddball, interhemispheric decreases were dominantly seen in the target response. For the DTF, the most apparent differences were in the alpha band. For the distractor response, these were decreases in intrahemispheric connections involving parietal electrodes, again with a preference for the right, which was consistent with the coherence. For the target, there was a large amount of significant inter- and intrahemispheric increases all originating from the same right-parietal region.

**Figure 3:**
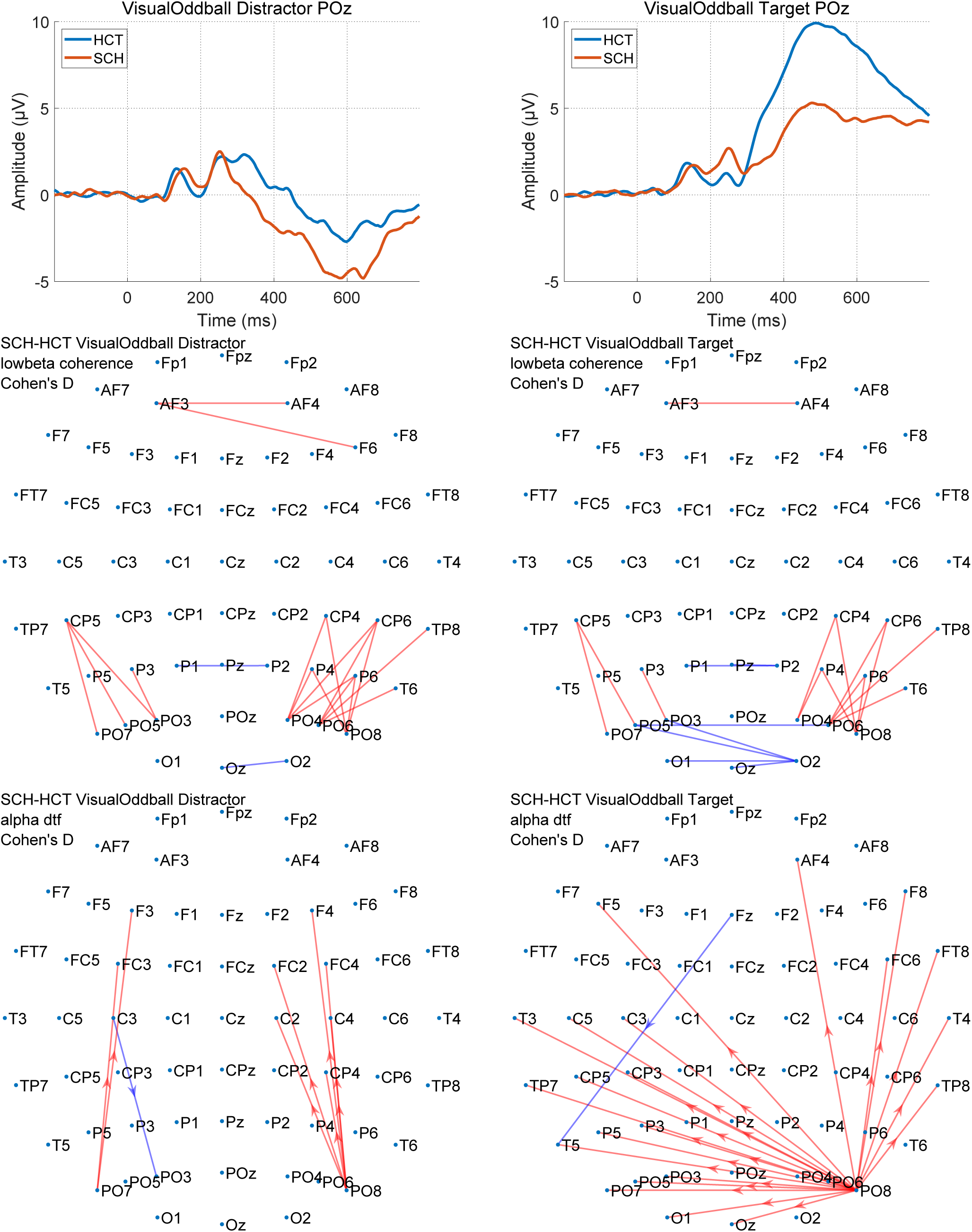
Visual oddball distractor and target.

### 3.5 Mismatch negativity

The coherence in high beta and the DTF in delta up to low beta did not pick up group differences. In the correlation and in all other bands of the coherence, we again found a pattern of intrahemispheric increases and interhemispheric decreases, which was most apparent in the alpha band (middle row of figure 4). Here too, these were parietal-occipitally located with increases located mostly in the right hemisphere. The long-range high-beta DTF decreases were mainly interhemispheric and the majority of the increases were intrahemispheric (bottom row of figure 4). Altered connections also almost exclusive involved electrodes in right-parietal/occipital regions.

**Figure 4:**
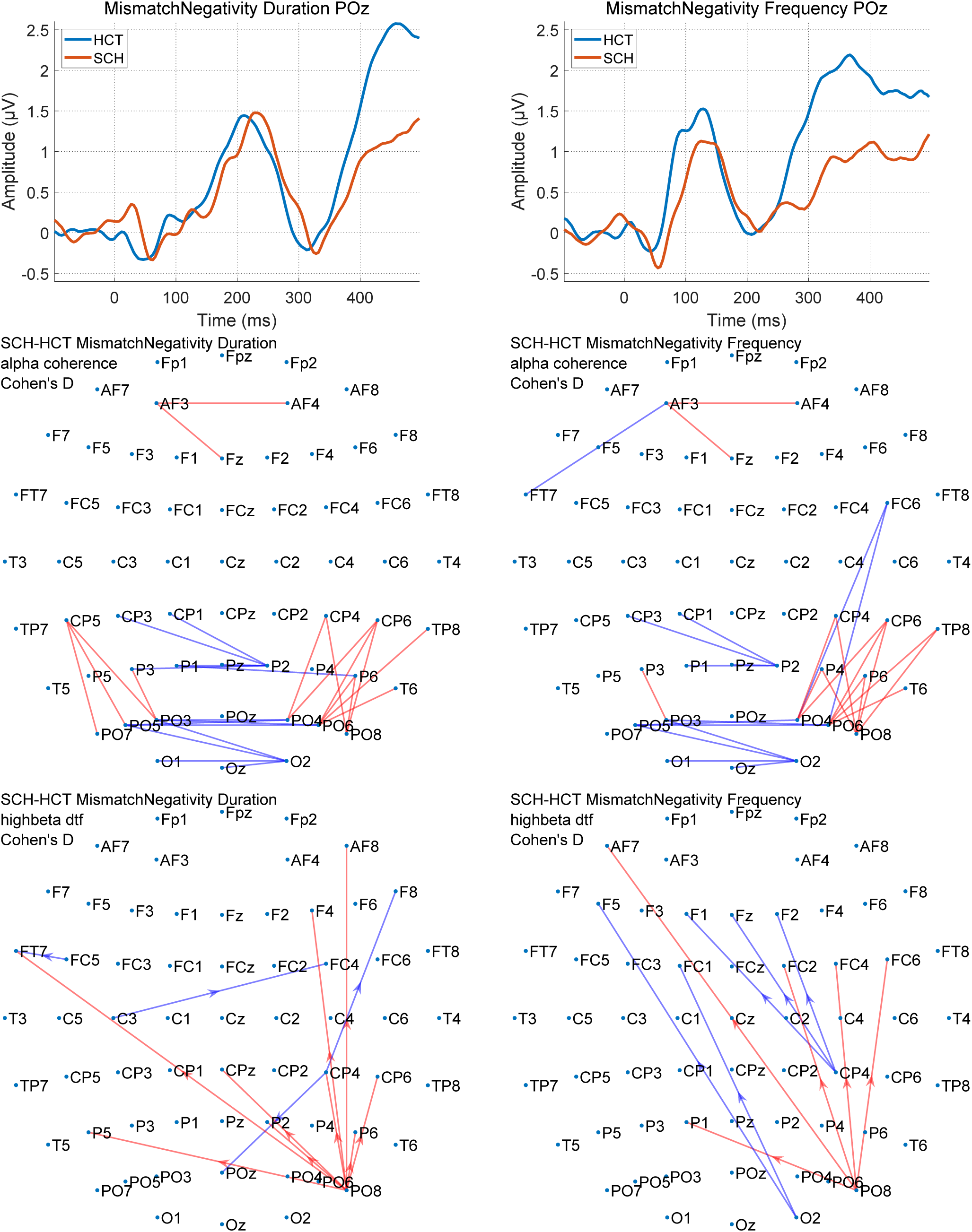
Mismatch negativity duration and frequency deviant.

## 4. Discussion

In this study, we wanted to verify whether network analysis on EEG sensor level can reveal the corpus callosum pathology in schizophrenia. Our main finding was that while we probed brain functioning with a range of paradigms, a very similar pattern of interhemispheric decreases and intrahemispheric increases in connectivity strength was revealed across paradigms, which was most apparent across the alpha and the low beta band. The DTF mostly found edge strength differences in long-range connections, while the coherence was better at detecting short-range connection differences. The correlation only picked up very few differences between the groups. In the mismatch paradigm, the found differences overlapped entirely with the differences identified by coherence method in low beta.

The inter- and intrahemispheric disruptions mainly occurred in parietal and occipital regions. Reduced connectivity between the hemispheres is believed to reflect the decreased processing capability that is seen in schizophrenia (Schlösser *et al.*, 2003; Hoptman *et al.*, 2012). This interhemispheric impairment is in line with the hypothesis of transcallosal disconnection (Crow, 1998), and the corpus callosum pathology found in schizophrenia (Foong *et al.*, 2000; Whitford *et al.*, 2010).

The pattern of increased intrahemispheric and decreased interhemispheric connectivity that we observed, is compatible with fMRI findings in 6 schizophrenic patients and six age and gender matched controls (Schlösser *et al.*, 2003). Schlosser et al observed volume reductions of the thalamus and altered connections between thalamus and cortex, leading to the interpretation that the pattern of enhanced intrahemispheric and reduced interhemispheric connectivity could be related to the disrupted ability to filter incoming thalamo-cortical information as proposed by (Jones, 1997). The increased intrahemispheric connectivity in our study might therefore reflect impaired integrative functions rather than increased efficiency.

Stimulus filtering or “sensory gating” has been known to be disrupted in schizophrenia (McDowd *et al.*, 1993; Javitt and Sweet, 2015). Schizophrenia patients have problems with suppression of active responses in the brain, which is in line with the already found weaker passive suppression of the P50, a peak at 50 milliseconds after stimulus that is normally suppressed when a stimulus is presented twice with a short time in between. In schizophrenia patients, the suppression of this second peak is less pronounced than in normal controls, showing their inability of stimulus filtering and hence a sensitivity to overstimulation (Braff, 1993).

This principle of sensory gating might extend to active suppression too (Gagnon *et al.*, 2000; Franke *et al.*, 2008; Chana *et al.*, 2013; Hayrynen *et al.*, 2016; Lavigne, Menon and Woodward, 2016; Bak *et al.*, 2017). When a deviant stimulus, or the absence of an expected stimulus, occurs in a train of frequent stimuli, the brain reacts to this occurrence in the form of the mismatch negativity, a negative peak around 200 milliseconds after the stimulus. In an active task with only one deviant stimulus type, a target stimulus to which the subject reacts with a button press, the brain eventually responds with a signal to the pyramidal pathway causing muscle movement. In an active task with both a target and a distractor as deviant stimuli, a new activity needs to occur in the brain: evaluation of the deviant stimulus. If the brain starts preparing to send a muscle movement signal while doing this evaluation, this preparation needs to be suppressed when a distractor is detected.

Lack of suppression of this preparation could explain why the three attention-related tasks are showing local connectivity increases, but not why this increase also occurs in rest. Since the order of the paradigms was not randomised, a possible explanation for why we see a similar, yet weaker local increase pattern at rest might lie in this sensitivity to overstimulation. It is extremely hard to eliminate every single *random* stimulus in a room filled with equipment. Any noise or light – coming from the physicist or the EEG equipment - can evoke a response in the brain, which could explain seeing slightly more increases during eyes open.

Across all four tasks, we found a very similar pattern of connectivity disturbance picked up by the coherence and the DTF. However, during the resting state, we found a large number of reductions in central and parieto-occipital electrodes influence in the DTF delta band on almost all electrodes. This was seen during eyes open and closed. Comparing schizophrenia to healthy controls at rest, Olejarczyk and Jernajczyk showed DTF influence originating from central, parietal and occipital electrodes towards all electrodes in both groups, and no influence from frontal (polar) or temporal electrodes (Olejarczyk and Jernajczyk, 2017). Although they did not show the differences in network strength, we believe that the DTF influence reductions we found for central and parieto-occipital regions are in line with their findings. Aside from these reductions, for rest with eyes open, we also found increases in the left hemisphere in temporal electrodes influencing frontal and parietal regions. These differences where not limited to the parietal region only, but the intrahemispheric nature of the increases seems in line with the coherence.

In our previous study, we extracted latencies and amplitudes (features) from the different stimulus responses (Laton *et al.*, 2014). We made a feature ranking based on significance, in which all but the last were significant after correction. Among those significant features, 14 originated from the auditory oddball, and only 3 from the visual oddball paradigm. Furthermore, using only auditory oddball features for classification generally yielded higher accuracies than visual oddball features. This had made us conclude that auditory evoked response peaks were the most predictive for schizophrenia. However, our current connectivity study revealed more significant effects in the visual than the auditory oddball paradigm, particularly more intrahemispheric increases. This indicates that neither of the analyses (peak versus connectivity) are able to capture the full endophenotype, so they should be considered complementary and not fully overlapping in information gain.

## 5. Conclusion

In this study, we could confirm that connectivity in the brain is disrupted in schizophrenia in a range of paradigms ranging from purely passive (rest with eyes closed) to active (oddball). We found indications that connectivity analyses complement traditional peak analyses in finding differences between schizophrenia and healthy people. Overall, the disruptions manifested as interhemispheric reductions and intrahemispheric increases, predominantly in parietal and occipital connections. We hypothesise that the interhemispheric reductions reflect transcallosal disconnection, while the intrahemispheric increases indicate the inability to suppress the response to incoming stimuli.

## Acknowledgement

We gratefully acknowledge the technical and nursing assistance of Mr Ignace Dewulf in obtaining the recordings for this study.

